# Synthesis of diagnostic quality cancer pathology images

**DOI:** 10.1101/2020.02.24.963553

**Authors:** Adrian B. Levine, Jason Peng, David Farnell, Mitchell Nursey, Yiping Wang, Julia R. Naso, Hezhen Ren, Hossein Farahani, Colin Chen, Derek Chiu, Aline Talhouk, Brandon Sheffield, Maziar Riazy, Philip P. Ip, Carlos Parra-Herran, Anne Mills, Naveena Singh, Basile Tessier-Cloutier, Taylor Salisbury, Jonathan Lee, Tim Salcudean, Steven J.M. Jones, David G. Huntsman, C. Blake Gilks, Stephen Yip, Ali Bashashati

## Abstract

Deep learning-based computer vision methods have recently made remarkable breakthroughs in the analysis and classification of cancer pathology images. However, there has been relatively little investigation of the utility of deep neural networks to synthesize medical images. In this study, we evaluated the efficacy of generative adversarial networks (GANs) to synthesize high resolution pathology images of ten histological types of cancer, including five cancer types from The Cancer Genome Atlas (TCGA) and the five major histological subtypes of ovarian carcinoma. The quality of these images was assessed using a comprehensive survey of board-certified pathologists (n = 9) and pathology trainees (n = 6). Our results show that the real and synthetic images are classified by histotype with comparable accuracies, and the synthetic images are visually indistinguishable from real images. Furthermore, we trained deep convolutional neural networks (CNNs) to diagnose the different cancer types and determined that the synthetic images perform as well as additional real images when used to supplement a small training set. These findings have important applications in proficiency testing of medical practitioners and quality assurance in clinical laboratories. Furthermore, training of computer-aided diagnostic systems can benefit from synthetic images where labeled datasets are limited (e.g., rare cancers). We have created a publicly available website where clinicians and researchers can attempt questions from the image survey at http://gan.aimlab.ca/.

## INTRODUCTION

Applications of deep learning to pathology images have shown great potential in a range of tasks, including identifying cancer,^1–3^ predicting patient outcomes,^4–6^ and classifying genomic driver mutations^7^ and molecular subgroups^8^ solely from images. These successes have exclusively used *discriminative* machine learning methods, which map data (such as images, text, or speech) to a class label, while *generative* methods, which model the underlying probability distribution of a data set and can synthesize examples of that data, have remained relatively unexplored. We were motivated to study generative modelling in pathology due to its numerous potential uses in education, clinical quality assurance, improving deep learning classifiers, and digital image processing (e.g., nuclei segmentation).

Generative adversarial networks (GANs) are a recently developed technique that have had great success in synthesizing high resolution, realistic images.^9^ GANs consist of a generator network (analogous to a counterfeiter), which creates synthetic images, and a discriminator network (the detector), which takes as input the synthetic images, as well as a set of real training images, and attempts to determine which are real and synthetic. The generator only receives as feedback whether the discriminator is fooled by the synthetic images and uses this to adjust its parameters; importantly, the generator does not at any point in training have direct access to the real images. Since the GAN concept was first published in 2014, there have been numerous advances in training methods, including the integration of deep convolutional architectures,^10^ and the development of the progressive GAN training method.^11^ Prior medical applications of GANs have included the synthesis of pathologic images of breast cancer,^12^ gliomas,^13^ and cervical dysplasia,^14^ as well as images of macular degeneration,^15^ dermatologic conditions,^16^ and several types of radiographic modalities.^17,18,19^ However, the images generated in these previous works were generally quite limited in size and low resolution.

There are several factors motivating our interest in generative modelling for medical images.^20^ Synthetically generated images would have significant educational value,^21^ in that they could provide a near endless source of novel examples of rare pathologies for teaching and proficiency testing, as well as avoiding confidentiality issues relating to including patient images in publicly available documents.^22^ Furthermore, these images could address a major issue in training deep neural networks for medical applications—the challenge in obtaining a sufficient amount of annotated training data for the model to capture high level features and prevent overfitting.^23^ Data compilation and annotation for medical applications typically requires the involvement of highly trained experts, and enriching data sets with synthetic images can leverage the work of these experts to maximize training material and decrease overall time and cost requirements.

We hypothesized that using GANs we could generate realistic-looking images that are adequately classifiable by pathologists, and can be utilized for educational purposes as well as leveraged to improve histopathology classifier performance, where limited datasets are available. Our results show that high quality tissue images as large as 1024 x 1024 pixels can be generated by GANs across five different cancer types as well as the five histotypes of ovarian carcinoma, and that expert pathologists cannot differentiate them from real images. Furthermore, we show that by leveraging this data, we can achieve performance improvements in deep learning models to classify cancer subtypes where limited data might be available. In aggregate, our work presents new opportunities in leveraging GANs to simulate realistic pathology images that can be utilized for educational and quality assurance purposes beyond their commonly advocated use for data augmentation in the machine learning community.

## RESULTS

### Cohort construction and generative adversarial network training

In order to represent a wide range of distinctive morphologic characteristics, 668 whole slide images (WSIs) corresponding to five cancer histotypes were taken from The Cancer Genome Atlas (TCGA) archive^24^ with the following breakdown: low-grade glioma (LGG; 333 slides), hepatocellular carcinoma (HCC; 78 slides), lung squamous cell carcinoma (LSCC; 72 slides), renal clear cell carcinoma (RCC; 104 slides), and papillary thyroid carcinoma (PTC; 81 slides). These five cancer types were selected based on their being relatively visually distinct and diagnosable based on a very small image. Furthermore, to show the utility of the methods in synthesizing pathology images for subtypes with subtle morphologic differences, we constructed a dataset of 354 WSIs corresponding to the five main histotypes of ovarian carcinoma from the BC Ovarian Cancer Research Program (OVCARE) with the following breakdown: high-grade serous (HGSC; 165 slides), low-grade serous (LGSC; 39 slides), endometrioid (ENC; 57 slides), clear cell (CCC; 56 slides), and mucinous (MUC; 37 slides). Following annotation of representative regions of tumor, smaller image patches of 1024×1024 pixels (at 40x optical zoom) were extracted from the WSIs and these were used to train various types of generative models.

### Comparison of different generative models suggests Progressive GANs as the most effective method of image generation

We compared the performance of various generative models including Progressive GANs, variational autoencoder ^25^, enhanced super resolution GAN (ESRGAN),^26^ and deep texture synthesis (see Supplemental Figure 4, Supplemental Methods 1).^27^ Upon visual inspection and quantitative comparison, we found that Progressive GANs were the only method that yielded high-resolution images (1024×1024 pixels) with low Frechet Inception Distance (FID) scores (lower scores correlate to better images), suggesting that high quality and visually recognizable images similar to real ones could be generated (Supplemental Table 6, Supplemental Methods 2). As such, we focused our efforts on the systematic evaluation of images generated by Progressive GANs. In addition, we experimented with using both individual Progressive GANs for each histological subtype of cancer and class-conditional GANs with multiple histological labels, and found that the GANs trained on a single subtype generated qualitatively more realistic images, with fewer artifactual distortions.

### Pathologist assessment of synthetic images reveals no major difference between real and synthetic images

Due to their lack of an objective performance function, it is challenging to directly evaluate the performance of GANs.^28^ A common test is to have human raters evaluate whether the images are real or synthetic. Using a survey consisting of a mixed set of real and synthetic images, we asked pathologists to classify each image as one of the five histotypes, rate whether they consider it to be of sufficient quality to enable histologic classification, and to determine if the image is real or synthetic. This is similar to the approach that was used previously for the assessment of retinal images.^15^

We created two surveys, the first of which was for the GANs trained on the TCGA slides of five different cancers. Four board-certified pathologists completed this survey (experience = 1-10 years). When asked to assign histotype, there was no statistically significant difference in the accuracy of the pathologists (compared to the true labels) when diagnoses were made from synthetic versus real images. In fact, pathologists performed slightly better on the synthetic images, with a median Cohen’s kappa of 0.77 (compared to the true diagnosis) for classifying the real images, compared to 0.78 for the synthetic images, supporting the equivalency of real and synthetic images (Fisher’s exact test *p = 1.0*; Table 1, Supplemental Table 2). The most challenging histotypes to diagnose were HCC and LSCC, which had substantially lower mean classification accuracies than the other three types (Supplemental Table 5, Supplemental Figure 6)). There was no major difference in the pathologist assessment of image quality between real and synthetic images (median 88% compared to 90% considered adequate for diagnosis, respectively; Fisher’s exact test *p = 0.82*). Finally, the pathologists did not demonstrate an ability to reliably distinguish the real from synthetic images, with a median performance of 54% (Fisher’s exact test *p = 1.0*), which is only slightly better than guessing with 50% chance.

**Table 1:**
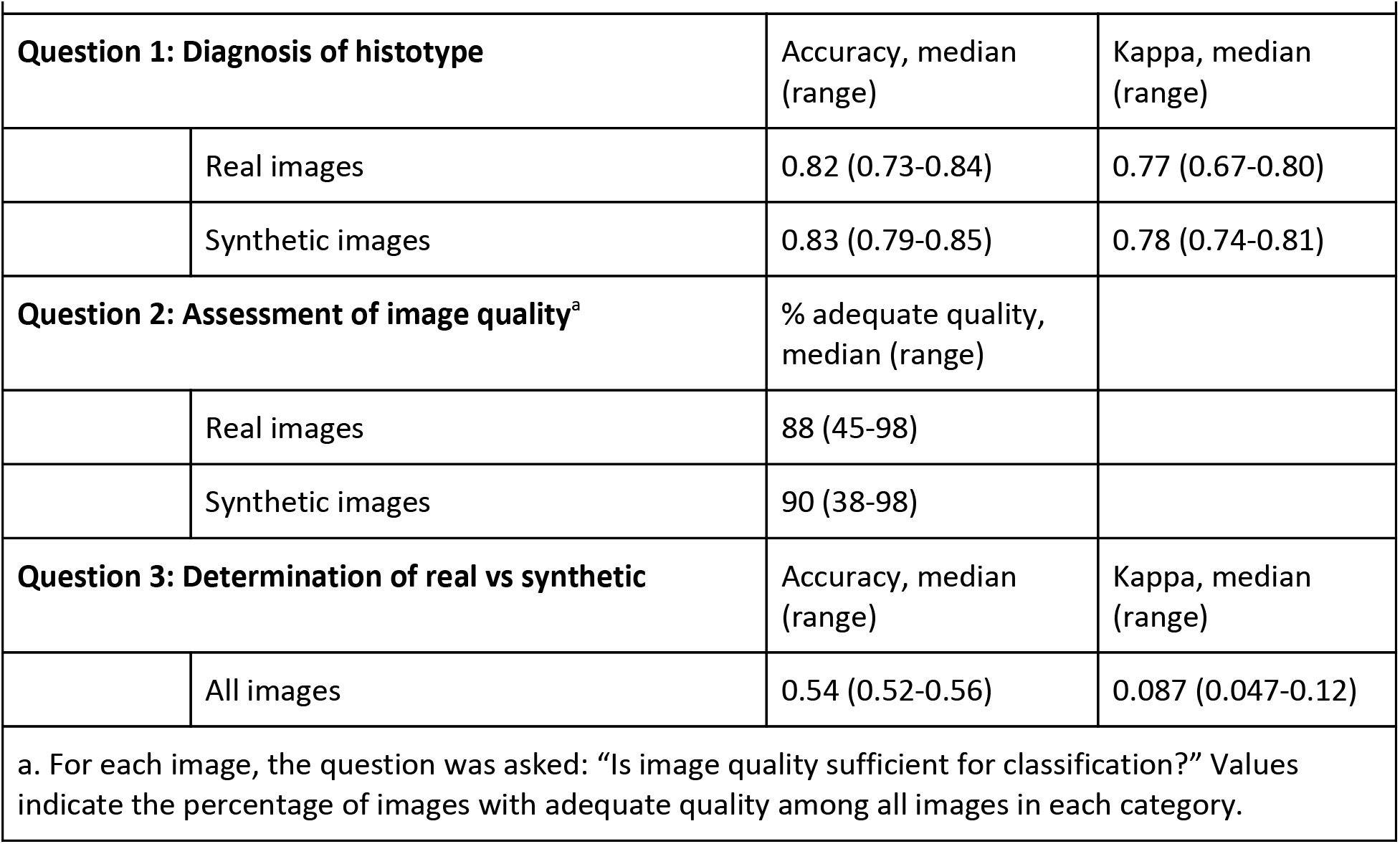
Pathologist assessment of TCGA image survey (n=4)

| **Table 1: Pathologist assessment of TCGA image survey (n=4)** |  |  |  |
| --- | --- | --- | --- |
| **Question 1: Diagnosis of histotype** |  | Accuracy, median (range) | Kappa, median (range) |
|  | Real images | 0.82 (0.73-0.84) | 0.77 (0.67-0.80) |
|  | Synthetic images | 0.83 (0.79-0.85) | 0.78 (0.74-0.81) |
| **Question 2: Assessment of image quality<sup>a</sup>** |  | % adequate quality, median (range) |  |
|  | Real images | 88 (45-98) |  |
|  | Synthetic images | 90 (38-98) |  |
| **Question 3: Determination of real vs synthetic** |  | Accuracy, median (range) | Kappa, median (range) |
|  | All images | 0.54 (0.52-0.56) | 0.087 (0.047-0.12) |
| a. For each image, the question was asked: “Is image quality sufficient for classification?” Values indicate the percentage of images with adequate quality among all images in each category. |  |  |  |

**Table 2:** Pathologist assessment of ovarian carcinoma image survey (n=5)

| **Table 2: Pathologist assessment of ovarian carcinoma image survey (n=5)** |  |  |  |
| --- | --- | --- | --- |
| **Question 1: Diagnosis of histotype** |  | Accuracy, median (range) | Kappa, median (range) |
|  | Real images | 0.73 (0.68-0.78) | 0.66 (0.60-0.73) |
|  | Synthetic images | 0.75 (0.65-0.85) | 0.69 (0.56-0.81) |
| **Question 2: Assessment of image quality<sup>a</sup>** |  | % adequate quality, median (range) |  |
|  | Real images | 81 (51-93) |  |
|  | Synthetic images | 91 (56-93) |  |
| **Question 3: Determination of real vs synthetic** |  | Accuracy, median (range) | Kappa, median (range) |
|  | All images | 0.54 (0.49-0.61) | 0.080 (-0.020-0.22) |
| a. For each image, the question was asked: "Is image quality sufficient for classification?" Values indicate the percentage of images with adequate quality among all images in each category. |  |  |  |

The second survey was for the GANs trained on the OVCARE slides, which included the five histotypes of ovarian carcinoma. Five board-certified pathologists in different geographic regions completed this survey (Table 2, Supplemental Table 3), all of whom had subspecialty expertise in gynecological pathology (experience = 5-30 years). For the first question of diagnosing the histotype, as expected the pathologists had lower overall accuracy than for the TCGA images but again performed slightly better on the synthetic images, with a median kappa of 0.66 compared to 0.69 for identifying the histotypes for the real and synthetic images, respectively (Fisher’s exact test *p = 1.0*). The most challenging histotypes to diagnose were ENC, HGSC, and LGSC, while a higher proportion of MUC and CCC were correctly diagnosed as such (Supplemental Table 5, Supplemental Figure 6). Overall evaluation of the image quality for histologic assessment revealed that the synthetic images had higher quality compared to real images (median 91% versus 81%; respectively), suggesting that GANs generated images that contain more histological distinctive features that aid the pathologist in better subtype identification. As in the first survey, the pathologists did not demonstrate an ability to reliably distinguish real from synthetic images, with a median accuracy of 54% (Fisher’s exact test *p = 0.88*).

Pathology trainees, ranging from beginner level (< 1 year pathology training) to senior resident/fellow also completed the OVCARE survey (see Supplemental Table 4 for results for each participant). Overall, these results were similar to those of the gynecological specialty pathologists, but with the expected lower classification accuracy for the histological subtypes. The trainees had a median classification kappa of 0.58 on the real images and 0.62 on the synthetic images, considered almost identical proportions of images to be high quality (median 90% of real images versus 92% of synthetic images), and had a median accuracy of 0.56 for distinguishing real from fake images. As expected, there was a trend towards better accuracy for classifying the histological subtypes for the more experienced trainees.

### Utility of synthetic data for training deep learning classifiers: achieving similar performance as real data

To further evaluate the quality of the synthetic images, as well as establish their value in building deep learning models for histotype classification where limited data is available, we evaluated whether the addition of synthetic images improved deep learning classifier accuracy similarly to the improvement that one would see with the addition of real images. We used a set of 52 additional ovarian cancer WSIs (17 HGSC, 10 CCC, 12 ENC, 6 MUC, 7 LGSC) that were not included in the training set of the GANs. We randomly split this data (10 times) into three sets of images for training, validation and testing. The resulting training set (referred to as *baseline training set*; mean number of patches in the set = 1,848) was then used to train a deep learning classifier to distinguish between the five ovarian cancer histotypes (see Methods). To evaluate whether the addition of GAN-generated synthetic images would improve machine learning classifier generalization similar to the addition of real images, we constructed two larger training sets with: (1) more real images added to the *baseline training set*, (2) synthetic images added to the *baseline training set* (mean number of images in each set = 42,098).

Using the three sets of training datasets, we then trained deep learning classifiers to distinguish between the five histotypes of ovarian cancer. While the median area under the curve (AUC) of the classifier for the baseline set across 10 runs was 0.840, the median AUC improved by adding more real or synthetic images to the baseline training set (*p*-value 0.099 and 0.0009, respectively; Supplemental Tables 8 and 9). More specifically, we achieved median AUC of 0.897 for the baseline augmented with real data and median AUC of 0.927 for the baseline augmented with synthetic images (Figure 3; Supplemental Table 8).

**Figure 1.**
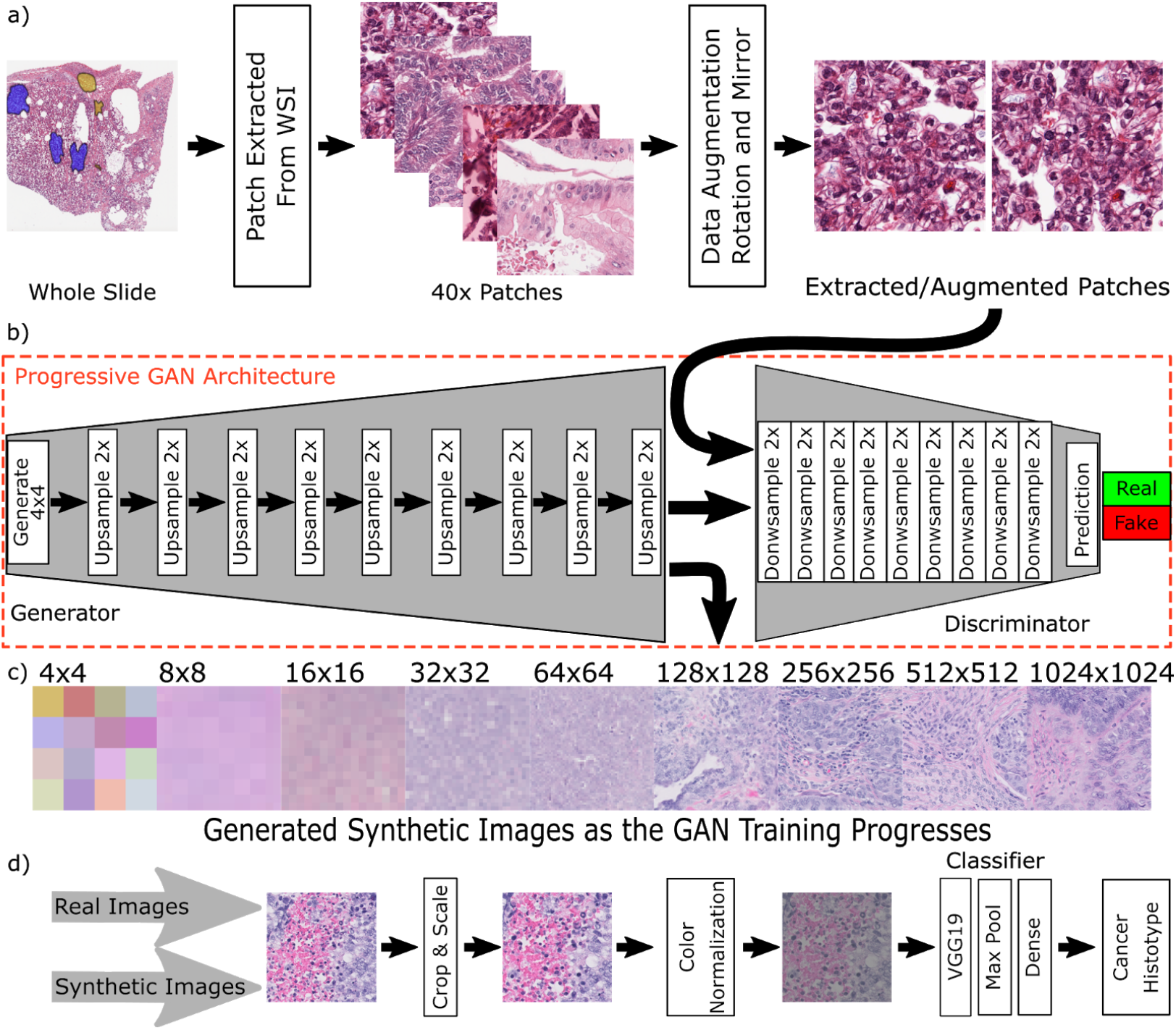
Workflow for Progressive GAN training. **a.** WSIs were annotated by pathologists and small image patches of size 1024 x 1024 were extracted and augmented with random rotation and mirroring. **b-c.** The image patches were used as the set of real images for training a Progressive GAN model, which begins with synthesizing very small images (4 x 4 pixels) and increases the size and resolution as training progresses. **d.** Following GAN training, synthetic images were generated and these, along with real training images, were preprocessed (through center cropping, scaling, and color normalization) and used to train a classifier to predict cancer histotype.

**Figure 2.**
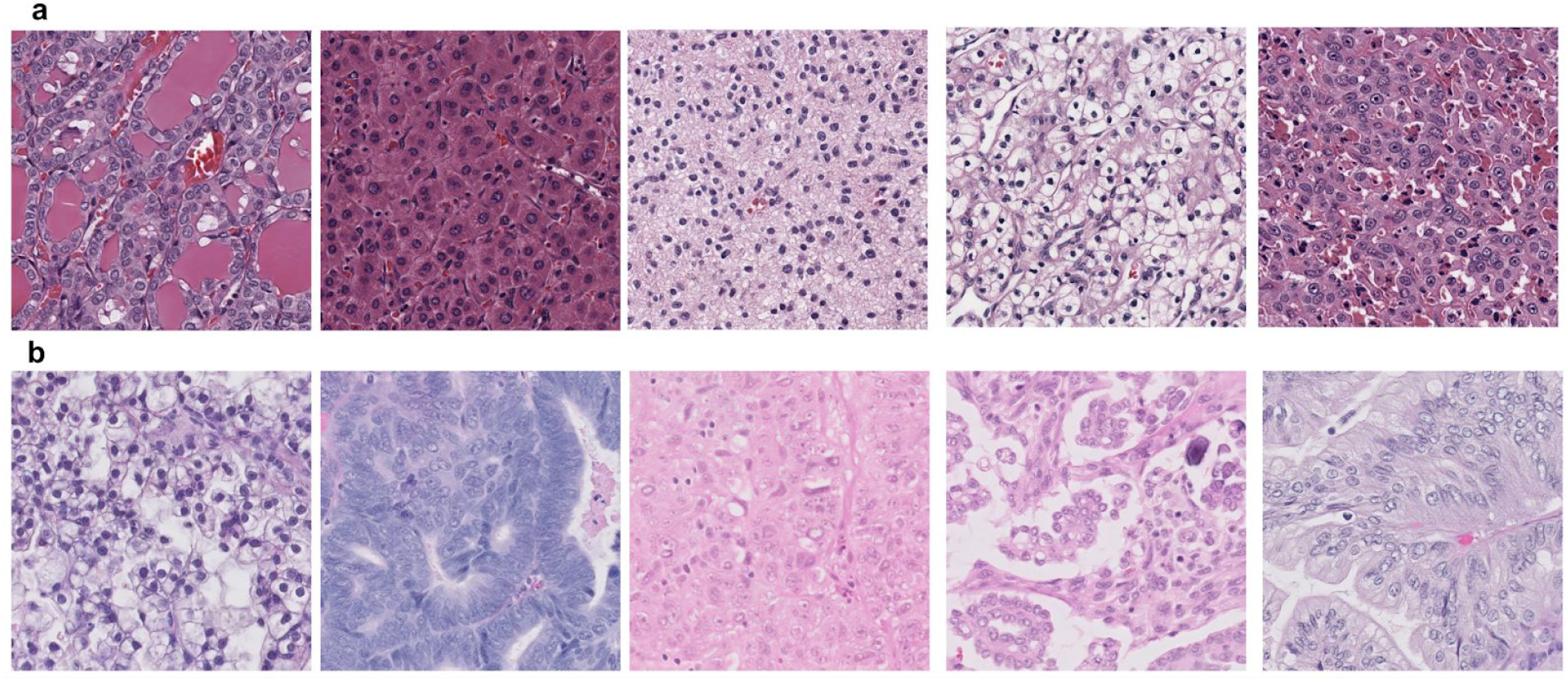
Representative examples of synthetic images. **a** Sample images (size=1024×1024 pixels) from GANs trained on TCGA Image Dataset (in order: PTC, HCC, LGG, RCC, SCC). **b** Sample images from GANs trained on OVCARE Dataset (in order: CCC, ENC, HGSC, LGSC, MUC)

**Figure 3.**
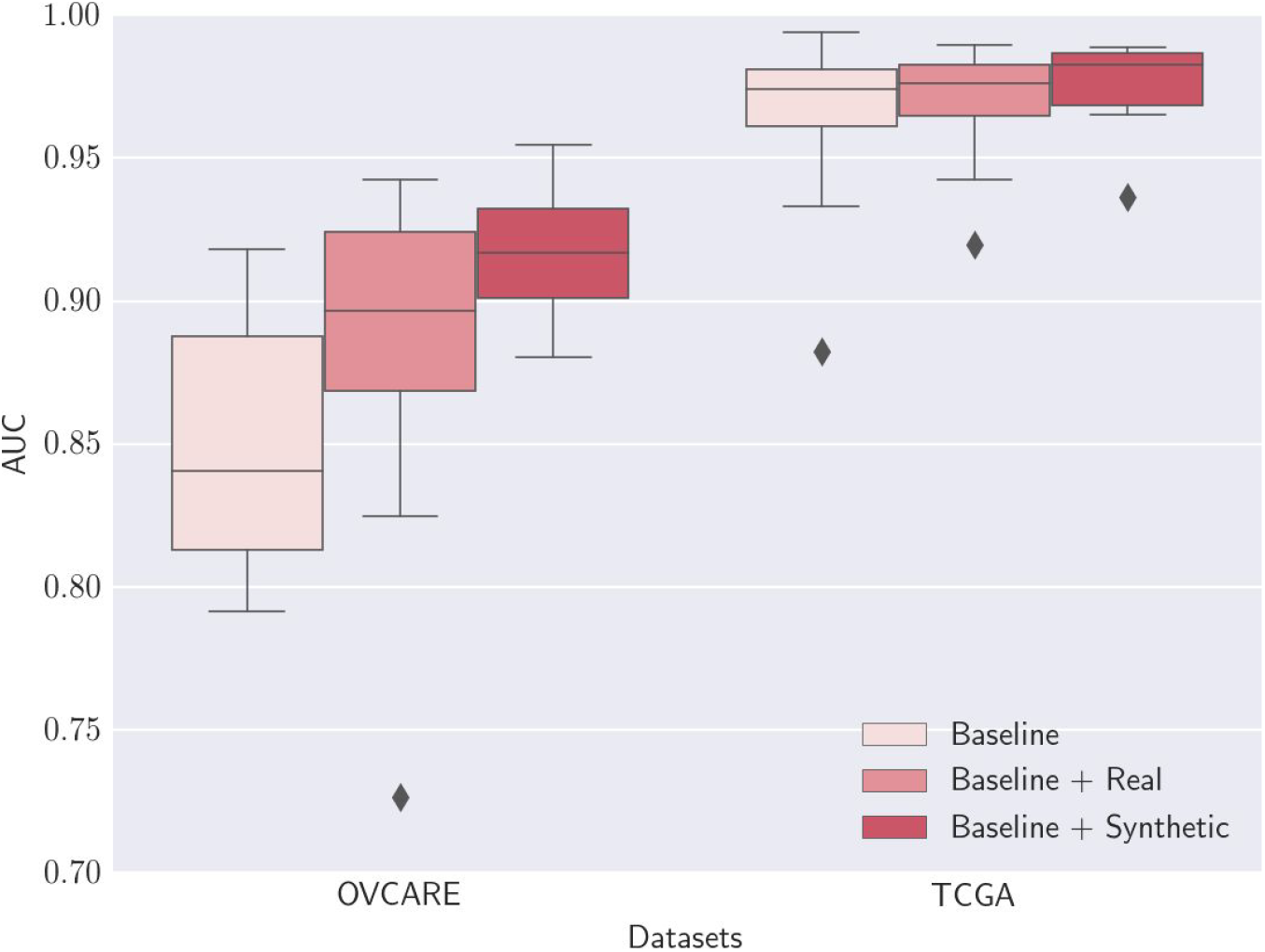
Results for classifiers trained on synthetic images. Multi-class area under the receiver operator curve for classifiers trained on the baseline training set, and training sets augmented with real images or with synthetic images. There was a statistically significant improvement in performance between the OVCARE baseline and baseline + synthetic image datasets (p=0.0009).

We repeated the same experiments with a set of additional 108 WSIs corresponding to various cancer histotypes (22 LGG, 20 HCC, 20 LSCC, 20 RCC, and 26 PTC) that were not included in the TCGA GAN training. While the median AUC of the classifier for the baseline set was 0.974, the performance improved by adding more real (median AUC = 0.976) or synthetic (median AUC = 0.983) images to the baseline training set (Figure 3; Supplemental Tables 8 and 9). While the improvement in performance for this data set was smaller than that for the ovarian carcinoma classifier, this is an easier classification task and the baseline performance was already very high, making it less likely that a large effect size could be observed.

It should be noted that the performance of the OVCARE and TCGA classifiers improved with the addition of either real or synthetic data to the baseline set. Similar to the assessment of image qualities by pathologists, although statistically not significant, augmentation with synthetic images led to more performance improvements compared to the addition of real images (Figure 3; Supplemental Tables 8 and 9). In aggregate, our results show that synthetic images generated by GANs perform at least similar to (if not better than) real images when used for training deep learning classifiers, suggesting that synthetic images have similar quality as real images.

## DISCUSSION

We present a flexible framework that can synthesize high quality histopathology images for a wide range of tissue types. To our knowledge these are the highest resolution synthetic pathology images that have been generated in the literature, and represent a significant advance in the use of generative deep learning in medicine. A set of pathologists from a range of subspecialties were able to classify the synthetic images based on a single very small patch, rated the image quality highly, and had no ability to reliably distinguish between real and synthetic images. We further demonstrate that the synthetic images are objectively useful as a form of data augmentation for classifier training, as enriching our training dataset with synthetic images improves the accuracy of neural networks trained to classify cancer histotypes. In comparison with other generative machine learning models, the GAN framework performed significantly better and generated qualitatively sharper and more realistic images. Future work is needed to increase the size and resolution of the generated images, as well as to investigate other technical variations within the GAN framework. Furthermore, with the increasing use of digital pathology in clinical practice^29^, generative models can be used for various digital image processing applications ^30,31,32^ and applied to forms of semi-supervised learning such as anomaly detection.^33^

Our use of two data sets demonstrates the breadth of input images that can be incorporated into our pipeline. The survey of images from GANs trained on TCGA slides was designed to be a straightforward classification task of five distinct cancer types, while the differentiation of the five histotypes of ovarian carcinoma is a challenging morphologic classification task for general pathologists, yet has evolved to become highly reproducible amongst experts.^34,35^ The accurate diagnosis of ovarian carcinoma is of critical importance in guiding therapy, as each ovarian carcinoma type has a distinct biomarker profile and natural history.^36,37^ The fact that the synthetic images could be reliably classified supports our assertion that they are of diagnostic quality, which in our view implies that they are usable for directly clinically relevant tasks. While the classification accuracy for ovarian carcinoma in our survey was lower than that reported in previous studies^35^, the task that we presented to the reference pathologists—the classification of cancer histotype based on a single small high-magnification image patch—was significantly more difficult than what is required in pathology practice, in which the whole slide or multiple slides are available for analysis.

Given the desire to use GANs to supplement datasets for rare tumors, an important factor to consider is the number of images that are required for GAN training. We have trained GANs using varying numbers of images, in part due to inherent limitations in the size of our training set and the need for manual annotation of the images. We found that overall, the minimum number of images needed to effectively train a GAN is approximately 8000, and that lower numbers than this result in an increased amount of artifact in the synthetic images. Despite the well described challenges in training GANs,^28,38^ we overall found that training proceeded very smoothly with the progressive GAN method, and very rarely experienced issues with training stability. In particular, we did not observe significant mode collapse, a common failure point in GAN training, in which the generator converges to output many copies of the same image. Some artifacts were encountered (see Supplemental Figure 3 for examples), including grid-like linear streaks in the images, lack of separation of nuclei in closely juxtaposed cells, in some cases forming linear chains of multiple nuclei, and, rarely and only with multi-label conditional GANs, images that were evenly split between two clearly different histotypes. Importantly, images rated of insufficient quality for interpretation were equally frequent amongst the synthetic and real images, demonstrating that significant artifacts were not generated at sufficient frequency to interfere with interpretation of the images.

Situated within the broader context of the implementation of artificial intelligence in the health care system, generative methods can address key concerns regarding data sharing and quality assurance.^39^ The current era of “big data” in medicine has the potential to significantly improve patient safety, but concurrently raises major privacy concerns, particularly in the context of data sharing between institutions and the possibility that combining multiple anonymized datasets can allow for re-identification.^22^ As the generator network in a GAN has no direct access to the patient images, these models, in combination with differential privacy strategies ^40^, can capture the features of a dataset yet be transferred with minimal risk to patient privacy.

While the initial studies applying deep learning to pathology used data sets of several hundred slides,^1,2^ more recent work has suggested that upwards of 10,000 slides may be needed to develop and validate a clinical support system that is capable of recognizing the full breadth of variability within pathology images.^41^ The resources necessary to assemble and securely store a dataset of this magnitude are a significant barrier for many institutions that would otherwise be interested in validating and implementing such systems, and GANs provide the ability to leverage available data and expand access to the technology. They would make it possible to run regular system quality assurance using synthetic images, at a fraction of the cost of manually curating cases to test the system, and ensure that the performance characteristics are acceptable day to day.

This same approach could be applied to proficiency testing of individual practitioners in morphology based specialities. It is well recognized that practitioners can fail to reach accepted standards of practice for a variety of reasons, including related to cognitive skills.^42^ While a number of strategies exist for physician performance assessment, they are inconsistently applied and may suffer from high cost and subjectivity (such as oral examinations or performance evaluations). Examinations to test diagnostic proficiency or cognitive ability, for example, are offered but the labor and cost associated with their production means they are only infrequently offered and then only to large cohorts. Proficiency testing on synthetic images could be done independent of a large cohort, with cases regularly refreshed, and at a fraction of the cost, with the output being objective and easily compared to the results of other practitioners. Such a system could also be used in education, to monitor progress through residency training.

In summary, generative adversarial networks can synthesize cancer pathology images that are indistinguishable from real images to expert observers and are useful as training data for other machine learning algorithms. The synthetic images have numerous applications, ranging from those that are directly and immediately clinically relevant, to others that are more research-focused and technical in nature. We are therefore optimistic that advances in generative modelling in medicine are an important step in the implementation of clinical artificial intelligence systems for improved efficiency and safety of care.

## METHODS

### Ethics statement

All experiments were conducted in accordance with the Declaration of Helsinki and the International Ethical Guidelines for Biomedical Research Involving Human Subjects. Anonymized archival tissue samples were retrieved from the pathology archive at the BC Cancer Ovarian Care Research Program (OVCARE), University of British Columbia and Vancouver General Hospital, and were digitized after approval by the institutional ethics boards.

### Data acquisition and processing

Whole slide images (WSIs) were acquired from The Cancer Genome Atlas Genomic Data Commons (GDC) Portal (https://portal.gdc.cancer.gov/).^24^ FFPE slides for ovarian cancer cases were acquired from the OVCARE archive and were scanned at 40x magnification on a Phillips IntelliSite Ultra Fast Scanner. Representative regions of tumor were annotated by either a board-certified pathologist, or a pathology resident under supervision by a board-certified pathologist. Following annotation, the WSIs were tessellated with stride=1024 (no overlap between patches) and smaller image patches of 1024×1024 pixels were extracted in TIFF format for training. During training, random horizontal mirroring was applied for further image augmentation.

Several types of cancer were used from the TCGA archive in order to represent a wide range of distinctive morphologic characteristics: low-grade glioma (333 slides; 71,043 image patches), liver hepatocellular carcinoma (78 slides; 59,894 images), lung squamous cell carcinoma (72 slides; 29,422 images), renal clear cell carcinoma (104 slides, 78,277 images), and papillary thyroid carcinoma (81 slides, 16,069 images). The slides used from the OVCARE archive included the five histotypes of ovarian carcinoma as follows: high-grade serous (165 slides; 75,861 images), low-grade serous (39 slides; 27,903 images), endometrioid (57 slides; 22,557 images), clear cell (56 slides; 28,246 images), and mucinous (37 slides; 19,000 images).

### Generative adversarial networks

For synthesizing histopathology images we trained a generative adversarial network on each of the histological subtype of cancers used. The images patches of 1024×1024 pixels were converted to the TFRecord format and used as the real image set in the discriminator training. The GAN training loop alternates between training the generator and the discriminator (with the other network’s parameters fixed), and attempts to solve the following two-player minimax game, with the value function given by *V*(*G,D*):

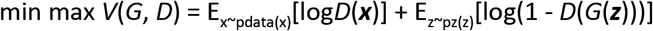

We used the Progressive GAN (ProGAN) training framework^11^, based on the openly available code from the authors’ implementation of this model (https://github.com/tkarras/progressive_growing_of_gans). The ProGAN training methodology begins with generating a very low resolution image (4 x 4 pixels) and adds new layers to the generator and discriminator to create higher resolution images with increased detail as training progresses. One of the fundamental challenges in GAN training is that higher resolution fake images are easily distinguished from real images. By beginning with very low resolution images, the ProGAN method improves training stability, with the additional benefit of reducing overall training time.

All codes were implemented in Python and used the Tensorflow deep learning library.^43^ Training was performed on 4 Nvidia Tesla V100 Graphical Processing Units (GPUs), and took approximately 3 days for each GAN. We did not alter the standard training parameters in the authors’ code. In brief, the networks were mainly composed of 3×3 convolutional layers with the leaky ReLU activation function, with upsampling and downsampling layers used in the generator and discriminator, respectively. In the discriminator, there was also a single mini-batch standard deviation layer inserted towards the end of the network, which helps to increase the variation in training data captured by the networks. Training was performed for a total of 12,000,000 images processed, with the batch size and learning rate adjusted based on the image resolution. The Adam optimizer was used^44^ with the initial learning rate=0.001 (this increased slightly at higher resolution images as per the standard training parameters), *β* =0, *β* =0.99, and *ε*=10^−8^. Following the completion of GAN training, the trained generator was used to synthesize images that were sent for pathologist review and used for data augmentation.

### Pathologist review

For the pathologist review and classification of synthetic images we created two online surveys using Google Forms (https://www.google.com/forms/about/)—one for the TCGA cancer types (low grade glioma, lung SCC, hepatocellular carcinoma, papillary thyroid carcinoma, and clear cell renal cell carcinoma) and one for the ovarian carcinoma subtypes (high-grade serous, low-grade serous, endometrioid, clear cell, and mucinous). The surveys included 150 synthetic and 150 real images and were evenly split between the five histotypes. The synthetic images were generated as consecutive images using the trained generator network, while the real images were selected randomly from the training images. In order to minimize the chance of bias, the images were not screened by the survey creators. Board-certified pathologists were tasked with reviewing each of the images and answering the following three questions about each:

1. What is the histotype?
2. Is the image of sufficient quality to enable classification?
3. Is the image real or synthetic?

For the TCGA cancer type survey, 4 pathologists (SY, DF, BS, MR) participated, from both academic and community practice settings, with 1-10 years of experience in practice. These pathologists had various areas of subspecialty expertise, including neuropathology, lung, renal, gastrointestinal, and molecular. For the ovarian cancer survey, 5 gynecologic subspecialty pathologists (BG, PI, CPH, AM, NS) participated, all from academic settings, and with 6-30 years of experience in practice.

### Deep learning-based histotype classification

For evaluating the utility of synthetic images as data augmentation, we trained deep learning classifiers to differentiate ovarian carcinoma subtypes as well as five cancer histotypes from TCGA. Each image classifier was trained and tested on an additional set of 52 (ovarian) and 108 (TCGA) WSIs that were not included in the training set of the GANs.

Image patches were pre-processed as follows: ovarian cancer images were centre-cropped to 768 x 768 pixels and down-sampled to 256 x 256, and TCGA images were down-sampled to 512 x 512. We then applied HSV colour jitter (during training only) with parameters adopted from ^45^ (brightness with a maximum delta of 0.25, contrast with a maximum delta of 0.75, saturation with a maximum delta of 0.25, and hue with a maximum delta of 0.04), and normalized to RGB pixel values between −1 and 1.

Using these images we randomly created ten splits of equal numbers of patients allocated to training/validation and testing data sets. The baseline training image set (average n = 1848 image patches for ovarian and n = 1967 for TCGA) was then augmented with either real images from the GAN training set or synthetic images generated by the GANs (n = 40,000 patches for ovarian and n = 2,000 for TCGA). Therefore, in total three different classifiers with identical hyperparameter settings were trained on each of the 10 patient splits, one for each of the following: baseline train images only (referred to as baseline setx), baseline images augmented with real images (referred to as baseline + real set), and baseline images augmented with synthetic images (referred to as baseline + synthetic set). On average, the validation and testing sets contained 7,850 and 18,802 image patches for ovarian and 14,617 and 12,994 image patches for TCGA, respectively.

The VGG19 network^46^ with batch normalization was chosen for the classifier based on its ease of use and robust performance. The PyTorch^47^ implementation was used with a modified last fully connected layer to classify the five classes (either five subtypes of ovarian carcinoma or five TCGA histotypes), with softmax activation to obtain the categorical distribution. We initialized each VGG19 model with weights trained on ImageNet, and all the weights in the model were optimized during back propagation. The Adam optimizer^44^ was used with learning rate=2*10^−4^, beta1=0.9, and beta2=0.999. Training was done with batch size=64 for 10 epochs and, using a single Tesla V100 GPU, took approximately 2 and 5 hours per model for the augmented TCGA and OVCARE datasets, respectively. Classification results are reported on the testing set using the weights of the VGG19 model taken from the training epoch with the highest validation accuracy. As the purpose was not to achieve the greatest absolute performance, but rather to evaluate the utility of synthetic data for improving performance, no significant hyperparameter tuning was performed.

### Statistical analysis

The sample size for the pathologist surveys was calculated using the work of Rotondi and Donner for an interobserver variability study.^48^ Specifically, the survey was powered to detect a difference of 0.1 between the kappa value for histological classification of real images versus the kappa value for classification of synthetic images. Supplemental Table 7 shows the minimum sample size for real and synthetic images for 4 and 5 pathologists. For more details regarding choice of sample size see Supplemental Methods 3.

### Data and source code availability

The data underlying the findings of this study are available from the corresponding author upon reasonable request. All TCGA digital slides are publicly available at the NCI Genomic Data Commons Portal (https://portal.gdc.cancer.gov/). Our source codes will be made freely available upon publication and can be applied to any tumor type.

## ACKNOWLEDGEMENTS

Eugene Shen, Jasleen Grewal, Sina Zareian, Dylan Lu, Davood Karimi

## REFERENCES

1. Ehteshami Bejnordi, B. et al. Diagnostic Assessment of Deep Learning Algorithms for Detection of Lymph Node Metastases in Women With Breast Cancer. JAMA 318, 2199–2210 (2017).

2. Steiner, D. F. et al. Impact of Deep Learning Assistance on the Histopathologic Review of Lymph Nodes for Metastatic Breast Cancer. Am. J. Surg. Pathol. 42, 1636–1646 (2018).

3. Chen, P.-H. C. et al. An augmented reality microscope with real-time artificial intelligence integration for cancer diagnosis. Nat. Med. 25, 1453–1457 (2019).

4. Mobadersany, P. et al. Predicting cancer outcomes from histology and genomics using convolutional networks. Proc. Natl. Acad. Sci. U. S. A. 115, E2970–E2979 (2018).

5. Courtiol, P. et al. Deep learning-based classification of mesothelioma improves prediction of patient outcome. Nat. Med. 25, 1519–1525 (2019).

6. Nagpal, K. et al. Development and validation of a deep learning algorithm for improving Gleason scoring of prostate cancer. NPJ Digit Med 2, 48 (2019).

7. Coudray, N. et al. Classification and mutation prediction from non-small cell lung cancer histopathology images using deep learning. Nat. Med. 24, 1559–1567 (2018).

8. Kather, J. N. et al. Deep learning can predict microsatellite instability directly from histology in gastrointestinal cancer. Nat. Med. 25, 1054–1056 (2019).

9. Goodfellow, I. J. et al. Generative Adversarial Networks. arXiv [stat.ML] (2014).

10. Radford, A., Metz, L. & Chintala, S. Unsupervised Representation Learning with Deep Convolutional Generative Adversarial Networks. arXiv [cs.LG] (2015).

11. Karras, T., Aila, T., Laine, S. & Lehtinen, J. Progressive Growing of GANs for Improved Quality, Stability, and Variation. in International Conference on Learning Representations (2018).

12. Quiros, A. C., Murray-Smith, R. & Yuan, K. Pathology GAN: Learning deep representations of cancer tissue. ArXiv (2019).

13. Hou, L. et al. Unsupervised Histopathology Image Synthesis. arXiv [cs.CV] (2017).

14. Xue, Y. et al. Synthetic Augmentation and Feature-based Filtering for Improved Cervical Histopathology Image Classification. arXiv [eess.IV] (2019).

15. Burlina, P. M., Joshi, N., Pacheco, K. D., Liu, T. Y. A. & Bressler, N. M. Assessment of Deep Generative Models for High-Resolution Synthetic Retinal Image Generation of Age-Related Macular Degeneration. JAMA Ophthalmol. 137, 258–264 (2019).

16. Ghorbani, A., Natarajan, V., Coz, D. & Liu, Y. DermGAN: Synthetic Generation of Clinical Skin Images with Pathology. arXiv [cs.CV] (2019).

17. Frid-Adar, M. et al. GAN-based Synthetic Medical Image Augmentation for increased CNN Performance in Liver Lesion Classification. arXiv [cs.CV] 321–331 (2018).

18. Han, C., Murao, K., Satoh, S.’ichi & Nakayama, H. Learning More with Less: GAN-based Medical Image Augmentation. arXiv [cs.CV] (2019).

19. Uzunova, H., Ehrhardt, J., Jacob, F., Frydrychowicz, A. & Handels, H. Multi-scale GANs for Memory-efficient Generation of High Resolution Medical Images. arXiv [eess.IV] (2019).

20. Yi, X., Walia, E. & Babyn, P. Generative adversarial network in medical imaging: A review. Med. Image Anal. 58, 101552 (2019).

21. McGaghie, W. C. Medical education research as translational science. Sci. Transl. Med. 2, 19cm8 (2010).

22. Price, W. N., 2nd & Cohen, I. G. Privacy in the age of medical big data. Nat. Med. 25, 37–43 (2019).

23. Levine, A. B. et al. Rise of the Machines: Advances in Deep Learning for Cancer Diagnosis. Trends Cancer Res. 5, 157–169 (2019).

24. Grossman, R. L. et al. Toward a Shared Vision for Cancer Genomic Data. N. Engl. J. Med. 375, 1109–1112 (2016).

25. Doersch, C. Tutorial on Variational Autoencoders. arXiv [stat.ML] (2016).

26. Wang, X. et al. ESRGAN: Enhanced Super-Resolution Generative Adversarial Networks. arXiv [cs.CV] (2018).

27. Gatys, L. A., Ecker, A. S. & Bethge, M. Texture Synthesis Using Convolutional Neural Networks. arXiv [cs.CV] (2015).

28. Salimans, T. et al. Improved techniques for training gans. in Advances in neural information processing systems 2234–2242 (papers.nips.cc, 2016).

29. Evans, A. J., Salama, M. E., Henricks, W. H. & Pantanowitz, L. Implementation of Whole Slide Imaging for Clinical Purposes: Issues to Consider From the Perspective of Early Adopters. Arch. Pathol. Lab. Med. 141, 944–959 (2017).

30. Mahmood, F. et al. Deep Adversarial Training for Multi-Organ Nuclei Segmentation in Histopathology Images. IEEE Trans. Med. Imaging (2019) doi:10.1109/TMI.2019.2927182.

31. Shaban, M. T., Baur, C., Navab, N. & Albarqouni, S. Staingan: Stain Style Transfer for Digital Histological Images. in 2019 IEEE 16th International Symposium on Biomedical Imaging (ISBI 2019) 953-956 (2019). doi:10.1109/ISBI.2019.8759152.

32. Xu, Z., Moro, C. F., Bozóky, B. & Zhang, Q. GAN-based Virtual Re-Staining: A Promising Solution for Whole Slide Image Analysis. arXiv [cs.CV] (2019).

33. Schlegl, T., Seeböck, P., Waldstein, S. M., Schmidt-Erfurth, U. & Langs, G. Unsupervised Anomaly Detection with Generative Adversarial Networks to Guide Marker Discovery. in Information Processing in Medical Imaging 146–157 (Springer International Publishing, 2017). doi:10.1007/978-3-319-59050-9_12.

34. Kommoss, S., Gilks, C. B., du Bois, A. & Kommoss, F. Ovarian carcinoma diagnosis: the clinical impact of 15 years of change. Br. J. Cancer 115, 993–999 (2016).

35. Köbel, M. et al. Diagnosis of ovarian carcinoma cell type is highly reproducible: a transcanadian study. Am. J. Surg. Pathol. 34, 984–993 (2010).

36. Köbel, M. et al. Ovarian carcinoma subtypes are different diseases: implications for biomarker studies. PLoS Med. 5, e232 (2008).

37. Köbel, M. et al. Biomarker-based ovarian carcinoma typing: a histologic investigation in the ovarian tumor tissue analysis consortium. Cancer Epidemiol. Biomarkers Prev. 22, 1677–1686 (2013).

38. Arjovsky, M. & Bottou, L. Towards Principled Methods for Training Generative Adversarial Networks. arXiv [stat.ML] (2017).

39. He, J. et al. The practical implementation of artificial intelligence technologies in medicine. Nat. Med. 25, 30–36 (2019).

40. Dwork, C. Differential Privacy: A Survey of Results. in Theory and Applications of Models of Computation 1–19 (Springer Berlin Heidelberg, 2008). doi:10.1007/978-3-540-79228-4_1.

41. Campanella, G. et al. Clinical-grade computational pathology using weakly supervised deep learning on whole slide images. Nat. Med. 25, 1301–1309 (2019).

42. Leape, L. L. & Fromson, J. A. Problem doctors: is there a system-level solution? Ann. Intern. Med. 144, 107–115 (2006).

43. Abadi, M. et al. TensorFlow: Large-scale machine learning on heterogeneous systems. (2015).

44. Kingma, D. P. & Ba, J. Adam: A Method for Stochastic Optimization. arXiv [cs.LG] (2014).

45. Liu, Y. et al. Detecting Cancer Metastases on Gigapixel Pathology Images. arXiv [cs.CV] (2017).

46. Simonyan, K. & Zisserman, A. Very Deep Convolutional Networks for Large-Scale Image Recognition. arXiv [cs.CV] (2014).

47. Paszke, A. et al. PyTorch: An Imperative Style, High-Performance Deep Learning Library. arXiv [cs.LG] (2019).

48. Rotondi, M. A. & Donner, A. A confidence interval approach to sample size estimation for interobserver agreement studies with multiple raters and outcomes. J. Clin. Epidemiol. 65, 778–784 (2012).

